# Cryo-EM structure of the RNA-rich plant mitochondrial ribosome

**DOI:** 10.1101/777342

**Authors:** Florent Waltz, Heddy Soufari, Anthony Bochler, Philippe Giegé, Yaser Hashem

## Abstract

The vast majority of eukaryotic cells contain mitochondria, essential powerhouses and metabolic hubs^1^. These organelles have a bacterial origin and were acquired during an early endosymbiosis event^2^. Mitochondria possess specialized gene expression systems composed of various molecular machines including the mitochondrial ribosomes (mitoribosomes). Mitoribosomes are in charge of translating the few essential mRNAs still encoded by mitochondrial genomes^3^. While chloroplast ribosomes strongly resemble those of bacteria^4,5^, mitoribosomes have diverged significantly during evolution and present strikingly different structures across eukaryotic species^6–10^. In contrast to animals and trypanosomatides, plants mitoribosomes have unusually expanded ribosomal RNAs and conserved the short 5S rRNA, which is usually missing in mitoribosomes^11^. We have previously characterized the composition of the plant mitoribosome^6^ revealing a dozen plant-specific proteins, in addition to the common conserved mitoribosomal proteins. In spite of the tremendous recent advances in the field, plant mitoribosomes remained elusive to high-resolution structural investigations, and the plant-specific ribosomal features of unknown structures. Here, we present a cryo-electron microscopy study of the plant 78S mitoribosome from cauliflower at near-atomic resolution. We show that most of the plant-specific ribosomal proteins are pentatricopeptide repeat proteins (PPR) that deeply interact with the plant-specific rRNA expansion segments. These additional rRNA segments and proteins reshape the overall structure of the plant mitochondrial ribosome, and we discuss their involvement in the membrane association and mRNA recruitment prior to translation initiation. Finally, our structure unveils an rRNA-constructive phase of mitoribosome evolution across eukaryotes.

---

Previously, we determined the full composition as well as the overall architecture of the *Arabidopsis thaliana* mitoribosome^6^. However, due to the difficulty to purify large amounts of *A. thaliana* mitoribosomes, mainly because of the low quantities of plant material usable for mitochondrial extraction, only a low-resolution cryo-EM reconstruction was derived. In order to obtain a high-resolution structure of the plant mitochondrial ribosome, we purified mitoribosome from a closely related specie, *Brassica oleracea var. botrytis*, or cauliflower (both Arabidopsis and cauliflower belong to the group of Brassicaceae plants), as previously described^6^(see Methods). We have recorded cryo-EM images for ribosomal complexes purified from two different sucrose gradient peaks (see Methods), corresponding to the small ribosomal subunit (SSU) and the full 78S mitoribosome. After extensive particle sorting (see Methods) we have obtained cryo-EM reconstructions for both types of complexes. The SSU reconstruction displayed an average resolution of 4.36Å (Extended Data Fig. 1). After multi-body refinement (3 bodies) and particle polishing in RELION3^12^ (see Methods), reconstructions were derived of the body of the SSU at 3.77Å, the head at 3.9Å and the head extension at 10.5 Å. The combined structure revealed the full plant mitoribosome SSU (Fig. 1a-d). As for the full mitoribosome, multi-body refinement of the LSU, SSU head and SSU body generated reconstructions at 3.50Å, 3.74Å and 3.66Å, respectively (Fig. 1e-g, Extended Data Fig. 1).

**Fig1.**
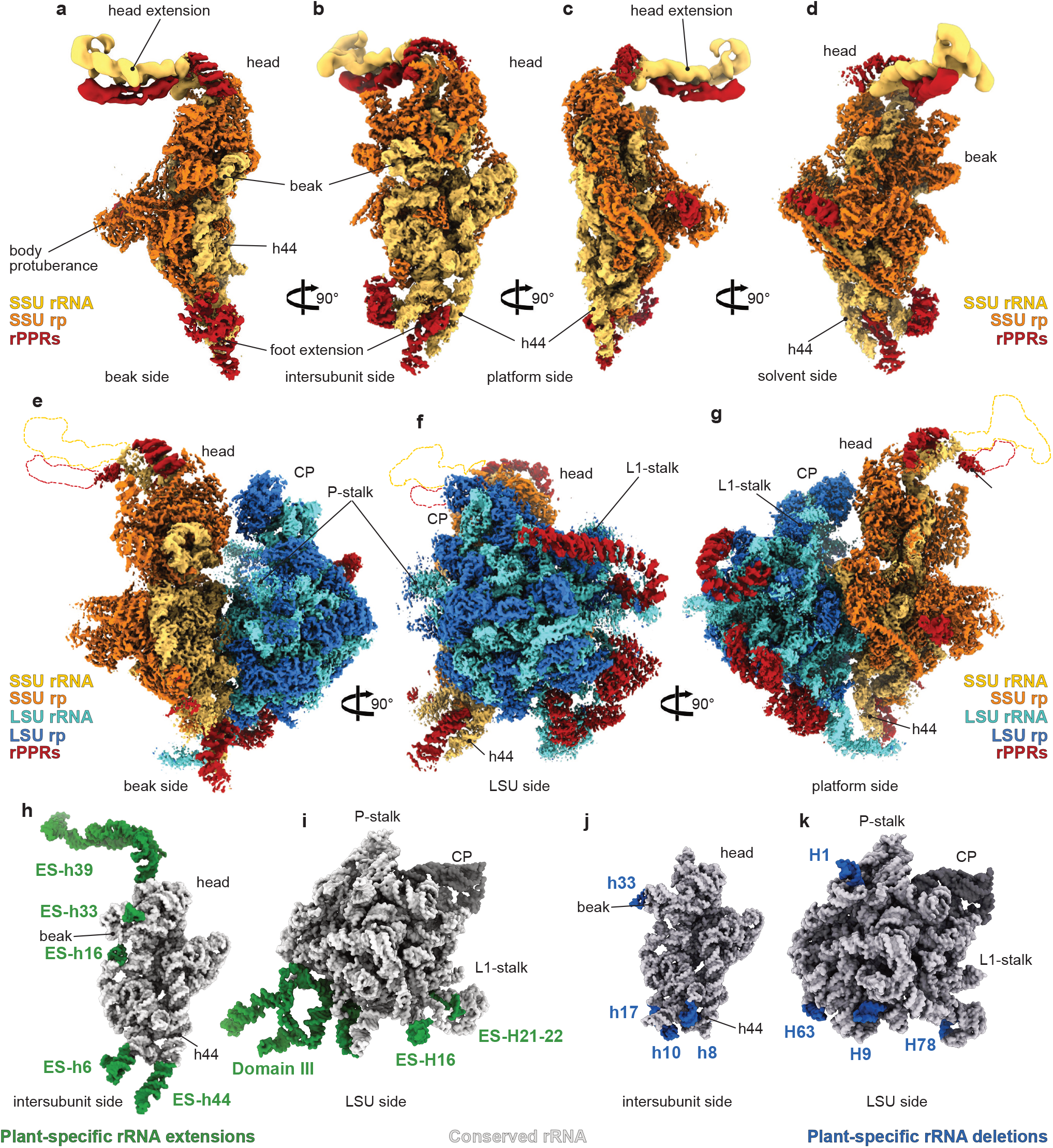
Overall structure of the plant mitochondrial ribosome. Composite cryo-EM map of the SSU alone (**a-d**) and the complete mitoribosome (**e-g**). rRNAs are colored in light blue (LSU) and yellow (SSU) and ribosomal proteins in blue (LSU) or orange (SSU). rPPR proteins are shown in red. **h-k** Arabidopsis mitoribosome rRNAs compared to E.coli ribosome. **h** and **j** comparison of the SSU, **i** and **k** comparison of the LSU, extensions in Arabidopsis rRNAs are shown in green, reductions are shown in blue. CP represents the central protuberance.

Density segmentations of our various cryo-EM reconstructions revealed the fine architecture of the plant mitoribosome showing a large rRNA core in interaction with numerous ribosomal proteins. Among those ribosomal proteins, 8 densities located at the surface of both subunits (3 on the LSU and 5 on the SSU) unambiguously display alpha-helical motifs characteristic of pentatricopeptide repeat proteins (Fig. 1a-g). Most of these ribosomal PPRs (rPPRs) are in direct interaction with large rRNA expansion segments (ESs) such as the head extension of the SSU (Fig. 1a-d). However, some of these SSU rPPRs appear to present a higher level of flexibility in the context of the full 78S, as they appear more scant compared to the SSU-only reconstruction, along with the ESs that they interact with. Consequently, we have focused our structural analysis of these SSU-rPPRs and their associated ESs on the SSU-only reconstruction.

Our cryo-EM reconstructions at near-atomic resolutions, along with our previous extensive MS/MS analysis^6^, allowed us to build a near-complete atomic models of the 78S mitoribosome. In contrast to its mammalian^10,13^ and trypanosoma^8^ counterparts, the plant mitoribosome is characterized by its largely expanded rRNAs, completely reshaping the overall structure of this mitoribosome. The 26S, 18S and 5S rRNAs are respectively 3,169, 1,935 and 118 nucleotides (nt) long, thus making the plant mitochondria SSU and LSU rRNAs 20% and 9% larger than their prokaryote counterparts, respectively^6^ (Fig. 1h-i). Nevertheless, while plant mitoribosomes contain more rRNA in general as compared to bacteria, they have lost a few rRNA helices present in bacteria (Fig. 1j-k).

As for the ribosomal proteins (45 in the LSU, 37 in the SSU), most of them are either universally conserved or mitochondria-specific constituting the common protein-core of almost all the known mitoribosomes, e.g. mS23, mS26 and mS29 on the SSU or mL46 and mL59/64 on the central protuberance of the LSU, thus confirming an acquisition of these proteins early during eukaryotes evolution.

The structure of the SSU revealed the exact nature of its several specific features, namely its large and elongated head additional domain, the body protuberance and its elongated foot (Fig. 1). The body protuberance is mainly formed by the mitoribosome-specific r-protein mS47, shared with yeast^7^ and trypanosoma^8^. Interestingly, mS47 is only absent in mammals, suggesting a loss of this protein during animal evolution. The body protuberance also contains one additional proteins (mS45) and extensions of uS4m (Extended Data Fig. 7), as well as additional protein densities. The foot of the small subunit is mainly reshaped by rRNA ESs and deletions, stabilized by plant-specific r-proteins, namely ribosomal PPR (rPPR) proteins. The SSU-characteristic helix 44 is 47 nt longer, thus slightly extending the SSU and forming part of the foot extension (Figs. 2 and 3e). This extension is stabilized by a rPPR protein itself connected to ES-h6 forming a three-way junction also stabilized by an additional rPPR protein. We could not identify these rPPRs with certainty based on their sequence because of the lack of resolution at these regions, however they can only correspond to rPPR1, 3a or 3b, identified in our previous work^6^, based on their number of repeats, they are here referred as rPPR*. In contrast, h8, h10 and h17 are reduced compared to their bacterial counterpart (Fig. 1j), leaving spaces for the plant specific additional proteins.

**Fig2.**
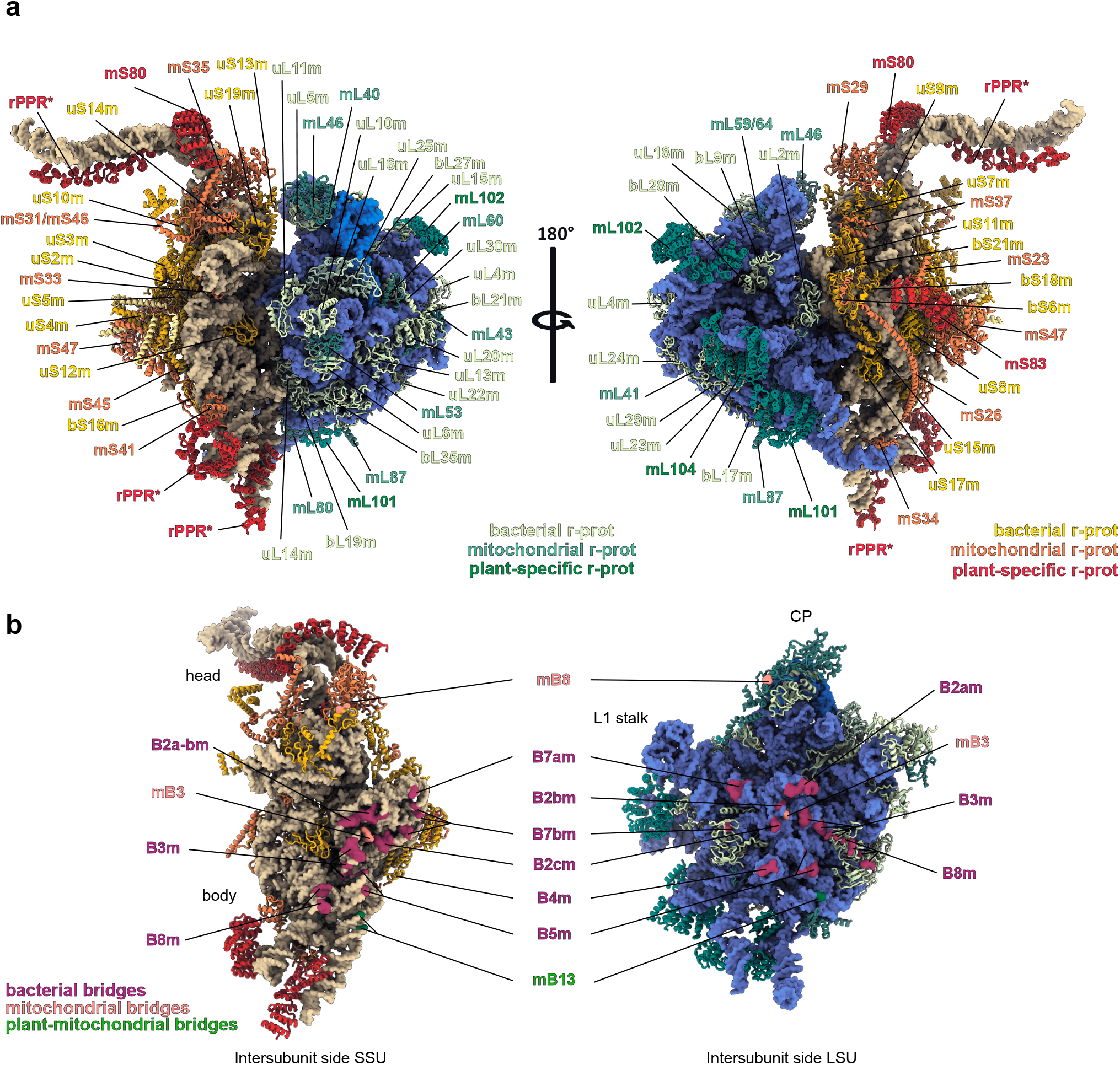
Atomic model of the plant mitoribosome. **a** The overall model of the plant mitoribosome with individual proteins annotated. rRNAs are colored in blue (LSU) and light brown (SSU) and ribosomal proteins in shades of blue and green (LSU) or shades of yellow and red (SSU), according to their conservation. **b** Intersubunit interfaces with conserved and novel observed intersubunit bridges highlighted. Bacterial ones are colored in purple, mitoribosomes ones in pink and plant ones in green.

**Fig3.**
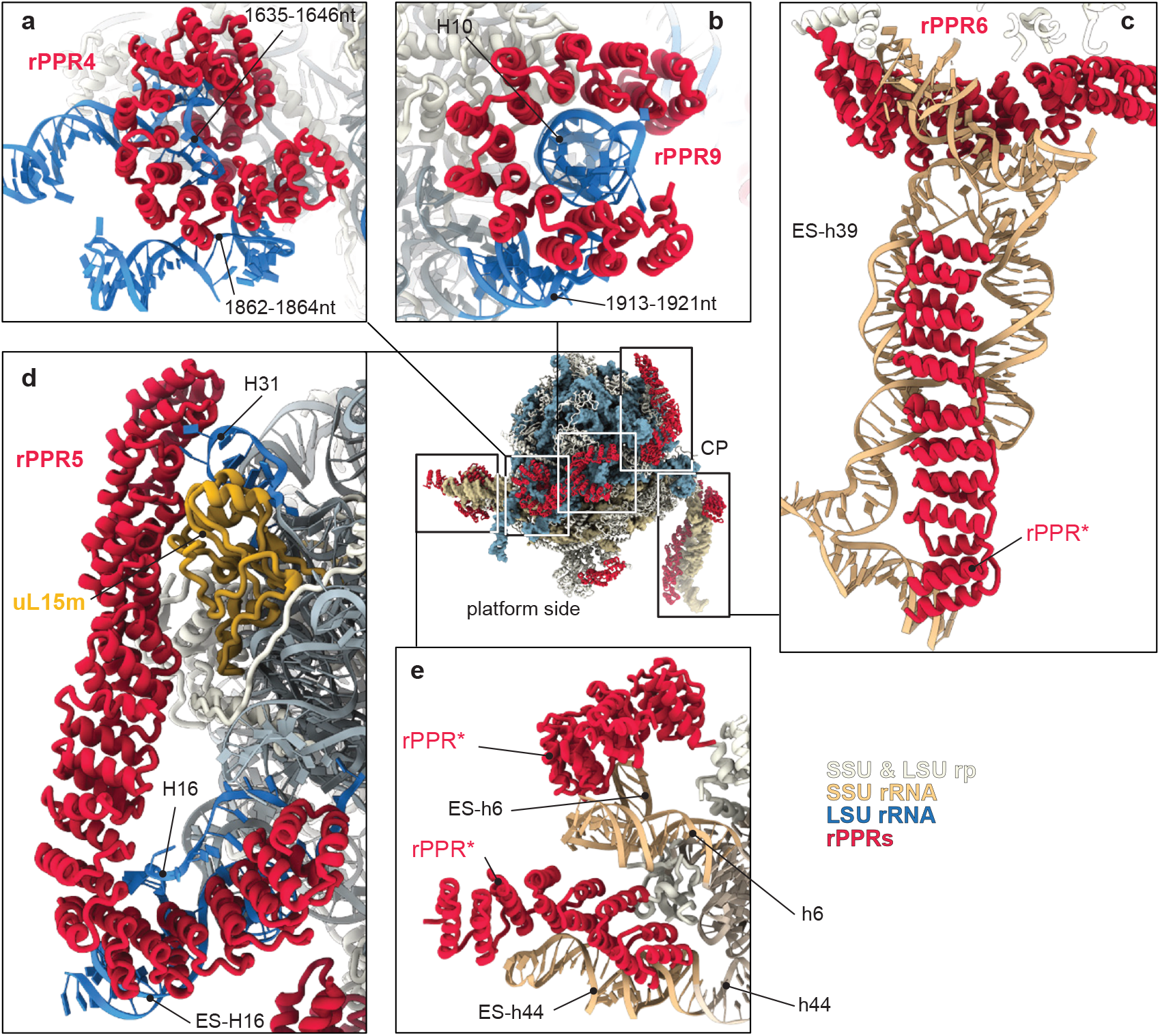
The plant mitoribosome PPR proteins. Overall view of the mitoribosome with PPR proteins highlighted in red, with zoomed-in view of each rPPR proteins (**a-e**) and their rRNA target and mode of binding. In **a** and **d** rPPR4 and rPPR5 stabilize single stranded segments, whereas in **b**, **c** and **e** rPPR proteins mainly interact with the backbone of the rRNAs.

Our analysis of the large SSU head extension revealed that it is indeed primarily shaped by a 370nt rRNA novel domain inserted in h39. Due to its high flexibility, it was refined with an average resolution around 10Å. Indeed, due to its movement relative to the head, but also the head movement relative to the body, the overall movement of the head extension is of large amplitude (~30°)(Extended Data Fig. 2), impairing the local resolution of this area. Nevertheless, the composition of the head extension can be determined unambiguously, as our data identifies secondary structure elements for both rRNA and rPPRs (Fig. 3c). It is mainly composed of rRNA, rooting from h39. From there, the extension forms a four-ways junction stabilized by a long PPR protein (rPPR6 or mS80), locking the whole additional domain in a position perpendicular to the intersubunit side. Past the four-ways junction, two of the rRNAs helices organize into two parallel segments, forming the core of the extension - one of the two helices ends in a three-way junction shaping the tip of the head-extension. The two parallel RNA helices are themselves contacted by a rPPR (Fig. 3c). However, local resolution is too low to clearly determine its exact identity (rPPR*). Interestingly, the protein bTHXm, previously identified by mass-spectrometry^6^ was found buried deep inside the small subunit head. This protein is only found in the plant mitoribosome as well as in chlororibosomes^4,5^ and ribosomes from the Thermus genus^14^.

On the back of the SSU, a large cleft extends the exit of the mRNA channel. This cleft is delimited by mS26, uS8m and h26 on one side and by mS47 and the rPPR mS83 on the other side (Fig. 4a). Similarly to all known PPRs, mS83 is predicted to be an RNA binder. In plants, the processes underlying the recruitment and correct positioning of mRNAs during translation initiation is unknown. Similar to other known mitochondrial translation systems, the Shine-Dalgarno (SD) and the anti-Shine-Dalgarno sequences are absent from both plant mRNAs and SSU rRNAs. Moreover, mRNAs have long 5’ untranslated regions (UTRs), similarly to yeast mitochondrial mRNAs^7^. Interestingly, half of the plant mitochondrial mRNA 5’UTRs harbors an A/purine rich sequence AxAAA located about 19nt upstream of the AUG (Extended Data Fig. 8). This distance correlates with the size of the extended mRNA exit channel and would put the purine-rich sequence in close vicinity to the rPPR protein mS83. We thus hypothesize that this plant-specific cleft may act as a recruitment platform for incoming mRNAs and / or additional factors. mS83 might recognize the AxAAA motif, thus recapitulating a SD/antiSD-like recognition system, using an RNA-protein interaction instead of an RNA-RNA interaction (Fig. 4a). An example of such possible rPPR-mediated initiation system may be found in mammalian mitochondria where the rPPR mS39 located on the SSU was proposed to accommodate the 5’UTR-less mRNAs from their 3’ end through a U-rich motif^15^. It is important to note that this A-rich cis-element is not found in all plant mitochondrial mRNAs, thus suggesting that this proposed mechanism for translation initiation would not be universal in plant mitochondria. Likewise, in chloroplasts, a third of the mRNAs do not possess SD-sequence, suggesting that at least two different mechanisms co-exist for translation initiation^16^.

**Fig4.**
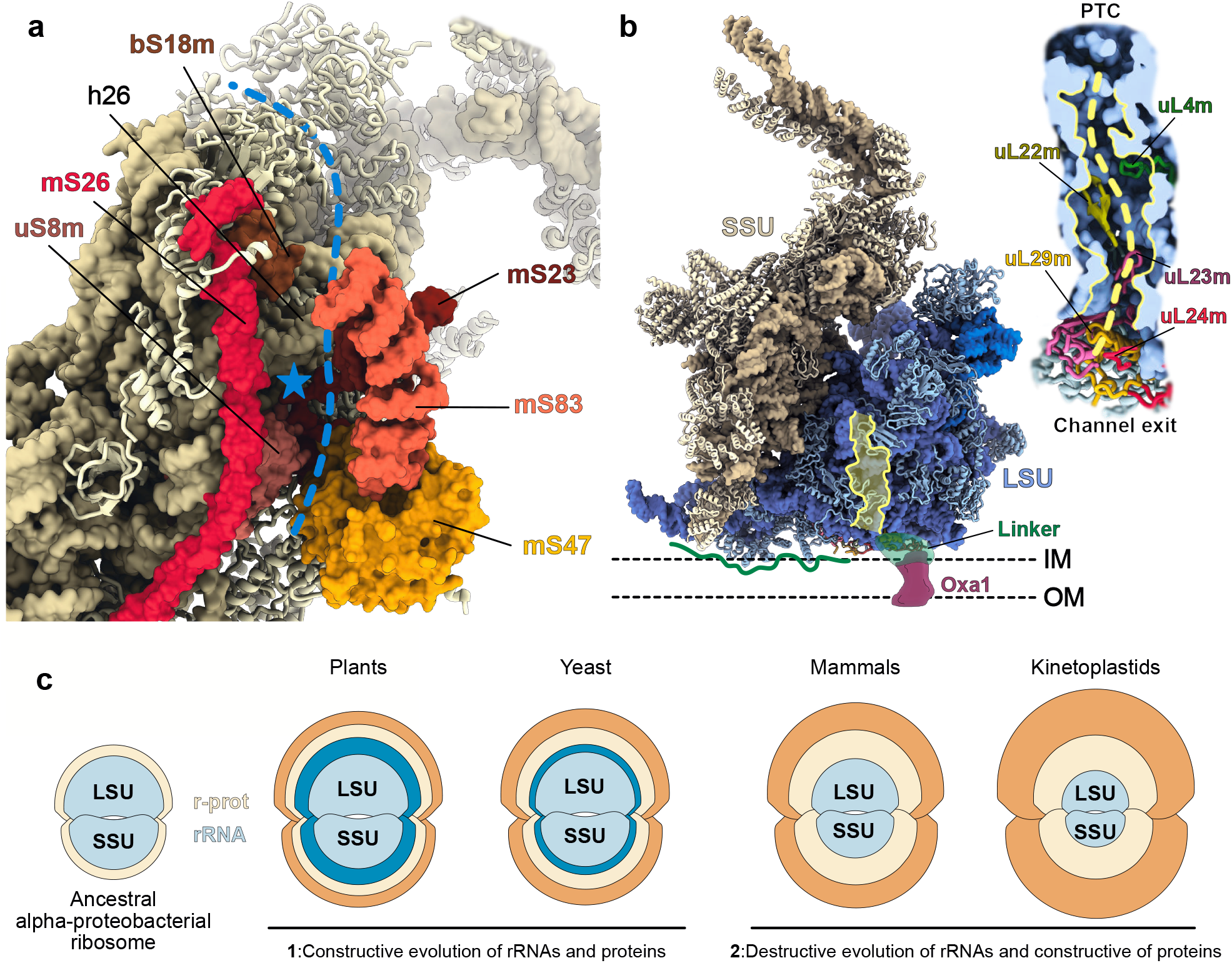
Specific features of the plant mitoribosome. **a** View of the SSU back channel and its proteins constituents. The possible mRNA path is highlighted by the dashed blue line and the blue star indicated the position of the AxAAA motif present in the 5’UTR of half of plant mitochondrial mRNAs. **b** Highlight of the plant mitoribosome peptide channel and possible mode of membrane attachment of the plant mitoribosome. Similar to yeast with Mba1^22^ plant mitoribosome would require an accessory protein factor (green linker) to tether the mitoribosome to the Oxa1 insertase. The green line represents the possible surface of interaction with the inner membrane (IM) of the mitochondria revealing an extensive surface mainly formed by the remodeled Domain III of the LSU. **c** Comparison of the different mitoribosomes described to date. Schemes of the ancestral alpha-proteobacterial is represented compared to the actual mitoribosomes. Light blue represent core bacterial rRNAs and yellow core bacterial r-proteins, blue and orange represent mitochondria specific rRNAs and r-proteins acquisition. Two main different evolutive paths have been taken by mitoribosomes, a mainly constructive one (**1**) represented by the yeast and plant mitoribosomes, where few proteins were lost and novel r-proteins were acquired as long as extended rRNAs, whereas in human and trypanosoma (**2**) rRNAs were highly reduced and more novel r-prot were acquired.

Similar to the SSU, the overall shape of the LSU is also strongly remodeled, even though a large portion of the core components are conserved (Fig. 2). Indeed, core functional components, such as the peptidyl-transferase center (PTC), the L7/L12 stalk and the central protuberance (CP) are similar to those found in bacteria. This conservation is in line with the globally bacterial-like intersubunit bridges (Fig. 2b). However, the LSU is reshaped by the conserved mitochondria-specific proteins e.g. mL41 or mL59/mL64 connecting the body of the LSU to the CP. Unlike all other mitoribosomes described to date, the plant mitoribosome CP includes a 5S rRNA. The core structure of the CP is quite conserved compared to prokaryotes. However, mitochondria-specific ribosomal proteins, namely mL40, mL59/64 and mL46, complement the classical bacterial-like CP and bind on top of the r-proteins bL27m, uL5m and uL18m. Interestingly, the same mitochondria-specific ribosomal proteins forming part of the plant CP are also present in other mitoribosomes even though they have lost their 5S rRNA, indicating that the acquisition of these mitoribosome-specific proteins occurred prior to the loss of the 5S in other mitoribosomes (Extended Data Fig. 5).

Interestingly, the universally conserved uL2 is split in two parts in the plant mitoribosome, its N-terminal part is encoded by a mitochondria encoded gene whereas its C-terminal part is encoded by a nucleus encoded gene, which could constitute a way of mito-nuclear crosstalk (Extended Data Fig. 7).

In yeast^7^, mammals^10,13^ and trypanosoma^8^ mitoribosomes, the peptide exit channel is highly remodeled by species-specific proteins (e.g mL45 or mL71). In plants however, the major part of the peptide channel and its exit are rather bacterial-like, with only minimal rearrangement of the surrounding rRNAs helices and a small extension of uL29m (Fig. 4b). In contrast to human and yeast^17^, a significant portion of the mitochondria encoded proteins are soluble proteins in plants (e.g. 7 soluble r-proteins are mitochondria encoded in *Arabidopsis*), thus it is conceivable that the plant mitoribosome does not systematically requires an association with the inner mitochondrial membrane, as it is the case for human and yeast^18,19^. However, it is likely that, at least in some cases, a non-ribosomal protein could link the mitoribosome to the mitochondrial insertase Oxa1. This hypothesis is supported by the observation that Oxa1 copurifies with immuno-precipitated plant mitoribosomes^6^(Fig. 4b).

The main plant-specific features of the LSU are the plant-specific proteins and rRNA ESs completely reshaping the back of the LSU, below the L1-stalk region. Indeed, running along the back of the LSU, from uL15m and H31 and contacting the largely extended and remodeled domain I rRNAs (ES-H21-22 and ES-H16-18), the 19-repeats rPPR protein mL102 (rPPR5) stabilizes these additional rRNAs extensions (Fig. 3d). Moreover, the domain III is extensively remodeled and holds several expansion segments, therefore helices of this domain were renamed pH53-59 (Fig. 1h, Extended Data Fig. 4). Indeed, this remodeled domain III has two main helices that largely extend in the solvent and are stabilized by two PPR proteins (rPPR4 and 9 or mL101 and mL104) (Fig 3a-b). These two rPPRs appear rigid and present numerous interactions with the rRNA. Thus, rPPR9 encapsulates the tip of helix H10, and stabilizes the end of pH59, one the newly formed helices of domain III. rPPR9 also directly contacts rPPR4 that wraps around a single stranded rRNA extension (1645-1644nt) and contacts the two major helices of domain III pH55 and pH57 (Fig. 3b). Interestingly, the rPPRs described here, along with those of the SSU, seem to hold a different mode of RNA binding as compared to the RNA recognition process of canonical PPR proteins. Indeed, PPR proteins usually bind ssRNA through a combinatorial recognition mechanism mainly involving two specific residues in each repeat (5 and 35) allowing each repeat to bind a specific nucleotide^20^, similar to other helical repeat modular proteins^21^. However, the structure obtained here revealed that several rPPR proteins bind the convex surface of double stranded rRNA, mainly through positively charged residues contacting the phosphate backbone of the rRNAs. This novel mode of RNA binding evidenced for rPPR proteins (Fig. 3) extends our understanding of the diversity of functions and modes of action held by PPR proteins. The position of this remodeled domain III on the full 78S suggests that rPPRs 4 and 9 might be involved in the attachment to the inner mitochondrial membrane, as they appear to strongly stabilize the structure of the whole domain (Fig. 4b).

In conclusion, the plant mitoribosome with its large rRNA ESs illustrates yet another route taken during mitoribosome evolution. It represents an augmented prokaryote-type ribosome. Based on a bacterial scaffold, it has both expanded rRNAs and an expanded set of proteins that were specifically recruited during eukaryote evolution in the plant clade. The structure of the plant mitoribosome could reflect the so-called “constructive phase” of mitoribosome evolution where both rRNAs and protein sets were augmented, in strong contrast with mammals and trypanosoma where rRNAs were considerably reduced^3^ (Fig. 4c). The structure presented here thus provides further insights into the evolution of mitoribosomes and the elaboration of independent new strategies to perform and regulate translation.

## METHODS

Methods and any associated references are available in the online version of the paper.

## ACKNOWLEDGEMENTS

This work has benefitted from the facilities and expertise of the Biophysical and Structural Chemistry platform (BPCS) at IECB, CNRS UMS3033, Inserm US001, University of Bordeaux. We thank A. Bezault for assistance with the Talos Arctica electron microscope. We thank L. Kuhn, J. Chicher and P. Hamman of the Strasbourg Espanade proteomic analysis for the proteomic analysis. We thank M. Sissler for her useful comments during the article redaction.

This work was supported by the “Centre National de la Recherche Scientifique”, the University of Strasbourg, by Agence Nationale de la Recherche (ANR) grants [MITRA, ANR-16-CE11-0024-02]] to PG and YH and by the LabEx consortium “MitoCross” in the frame of the French National Program “Investissement d’Avenir” [ANR-11-LABX-0057_MITOCROSS, as well as by a European Research Council Starting Grant (TransTryp ID:759120) to YH.

## DATA AVAILABILITY

The cryo-EM maps of the mitoribosome have been deposited at the Electron Microscopy Data Bank (EMDB) with accession codes XXXX. Corresponding atomic models have been deposited in the Protein Data Bank (PDB) with PDB codes XXXX.

## AUTHOR CONTRIBUTIONS

PG, FW, and YH designed and coordinated the experiments. FW purified the mitochondria and mitochondrial ribosomes. HS acquired the cryo-EM data. HS and YH processed the cryo-EM results. HS, AB and FW built the atomic models. FW, HS and YH interpreted the structure. PG, FW, HS and YH wrote and edited the manuscript.

## ONLINE METHODS

### Mitochondrial ribosome purification

Cauliflower (*Brassica oleracea* var. botrytis) was used here as it is best suited for large scale biochemical, structural analyses as compared to Arabidopsis. Cauliflower belongs to the same family of plants as Arabidopsis, Brassicaceae, thus making it an optimal model for this study, allowing the preparation of large quantities of highly pure mitochondria. However, the genome of Cauliflower is not sequenced, and the closest fully sequenced member of the family (*Brassica oleracea* subsp. oleracea) is poorly annotated. Still, protein sequence identities between members of the Brassicaceae family are higher than 90%, thus facilitating proteomics identification of cauliflower proteins. Combining the proteomics results from Waltz 2019 and those obtained in this study it is evident that no difference in terms of protein composition could be observed between Cauliflower and Arabidopsis. Hence, to facilitate comprehension and analysis we positioned Arabidopsis proteins in the cauliflower map.

For the mitochondria purification, fresh cauliflower inflorescence tissue was blended in extraction buffer containing 0.3 M mannitol, 30 mM sodium pyrophosphate (10.H_2_O), 0.5 % BSA, 0.8 % (w/v) polyvinylpyrrolidone-25, 2mM beta-mercaptoethanol, 1 mM EDTA, 20 mM ascorbate and 5 mM cysteine, pH 7.5. Lysate was filtered and clarified by centrifugation at 1.500 g, 10 min at 4°C. Supernatant was kept and centrifuged at 18.000 g, 15 min at 4°C. Organelle pellet was re-suspended in wash buffer (0.3 M mannitol and 10 mM phosphate buffer, 1 mM EDTA, pH 7.5) and the precedent centrifugations were repeated once. The resulting organelle pellet was re-suspended in wash buffer and loaded on a single-step 30 % Percoll gradient (in wash buffer without EDTA) and run for 1h30 at 40.000 g. Mitochondria are were retrieved, washed two times before being flash frozen in liquid nitrogen.

For mitoribosome purification, mitochondria were re-suspended in Lysis buffer (20 mM HEPES-KOH, pH 7.6, 100 mM KCl, 30 mM MgCl_2_, 1 mM DTT, 1.6 % Triton X-100, 100 μg/mL chloramphenicol, supplemented with proteases inhibitors (Complete EDTA-free)) to a concentration of 1 mg/mL and incubated for 15 min in 4°C. Lysate was clarified by centrifugation at 30.000 g, 20 min at 4°C. The supernatant was loaded on a 40% sucrose cushion in Monosome buffer (same as lysis buffer without Triton X-100 and 50 μg/mL chloramphenicol) and centrifuged at 235.000 g, 3h, 4°C. The crude ribosomes pellet was re-suspended in Monosome buffer and loaded on a 10-30 % sucrose gradient in the same buffer and run for 16 h at 65,000 g. Fractions corresponding to mitoribosomes were collected, pelleted and re-suspended in Monosome buffer.

### Grid preparation

4 μL of the samples at a concentration of 2 μg/μl was applied onto Quantifoil R2/2 300-mesh holey carbon grid, which had been coated with thin home-made continuous carbon film and glow-discharged. The sample was incubated on the grid for 30 sec and then blotted with filter paper for 2.5 sec in a temperature and humidity controlled Vitrobot Mark IV (T = 4°C, humidity 100%, blot force 5) followed by vitrification in liquid ethane pre-cooled by liquid nitrogen.

### Single particle cryo-electron microscopy data collection

For the two data-sets (Full and dissociated complexes), data collection was performed on a Talos Artica instrument (FEI Company) at 200 kV using the EPU software (FEI Company) for automated data acquisition. Data were collected at a nominal underfocus of −0.5 to −2.7 μm at a magnification of 120,000 X yielding a pixel size of 1.21 Å. Micrographs were recorded as movie stack on a Falcon II direct electron detector (FEI Compagny), each movie stack were fractionated into 20 frames for a total exposure of 1 sec corresponding to an electron dose of 60 ē/Å2.

### Electron microscopy image processing

Drift and gain correction and dose weighting were performed using MotionCor2^25^. A dose weighted average image of the whole stack was used to determine the contrast transfer function with the software Gctf^26^. The following process has been achieved using RELION 3.0^27^. Particles were picked using a Laplacian of gaussian function (min diameter 260 Å, max diameter 460 Å). For the full mitoribosome, after 2D classification, 153,608 particles were extracted with a box size of 400 pixels and binned four fold for 3D classification into 10 classes. Four classes depicting high-resolution features have been selected for refinement. The complex has been focused refined with a mask on the LSU, the body and the head of the SSU, yielding respectively 3.50, 3.66 and 3.74 Å resolution. Determination of the local resolution of the final density map was performed using ResMap^28^.

For the dissociated subunits, following a 2D classification, 132,130 particles for the LSU and 120,350 particles for the SSU were extracted with a box size of 400 pixels and binned four fold for 3D classification into 8 classes for each subunits. Five subclasses depicting high-resolution features have been selected for the SSU refinement with 73,670 particles. After Bayesian polishing a multi-body refinement has been performed using mask on the body, the head and the RNA expansion on the head, yielding respectively 3.77, 3.9 and 10.5 Å resolution. Three classes have been selected for the LSU refinement yielding a resolution of 3.96 Å. Determination of the local resolution of the final density map was performed using ResMap^28^.

### Structure building and model refinement

The atomic model of the plant mitoribosome was built into the high-resolution maps using Coot, Phoenix and Chimera. Atomic models from *E.coli* ribosome (5kcr)^29^, yeast mitoribosome (5mrc)^7^, human mitoribosome (6gaw)^15^ and trypanosoma mitoribosome (6hiv)^8^ were used as starting points for protein identification and modelisation. The online SWISS-MODEL service was used to generate initial models for bacterial and mitochondria conserved r-proteins. Models were then rigid body fitted to the density in Chimera^30^ and all subsequent modeling was done in Coot^31^.

For the LSU and SSU ribosomal RNA, the 16S and 23S from *E.coli* were docked into the maps and used as templates from positioning and reconstruction. A multiple sequence alignment of several plant mitochondrial ribosomes and *E.coli* ribosome was performed in order to determine the additional or depleted domains of *A.thaliana* mitochondrial ribosome. Ponctual differences were done in Chimera using the “swapna” command line.

In order to build the additional rRNA domains of *A.thaliana*, the co-variation algorithm LocARNA webservice (http://rna.informatik.uni-freiburg.de) was used to determine secondary structure of these domains. At that point, the secondary structure prediction was used to build the 3D model in Chimera using the “build structure” tools followed by manual adjustments.

Proteins with clear homologs in either mammalian or yeast mitoribosomes as long as *E.coli* ribosome were build using Phyre2 and SWISS-Model.

The global atomic model was refined with VMD using the Molecular Dynamic Flexible Fitting (MDFF) then with PHENIX using a combination of real and reciprocal space refinement for proteins and ERRASER for RNA.

### Proteomic and statistical analyses of mitochondrial ribosome composition

Mass spectrometry analyses of the ribosome fractions were performed at the Strasbourg-Esplanade proteomic platform and performed as previously^6^. In brief, proteins were trypsin digested, mass spectrometry analyses and quantitative proteomics were carried out by nano LC-ESI-MS/MS analysis on AB Sciex TripleTOF mass spectrometers and quantitative label-free analysis was performed through in-house bioinformatics pipelines.

### Figure preparation

Figures featuring cryo-EM densities as well as atomic models were visualized with UCSF ChimeraX^32^.

---

**Extended Data Fig. 1.**
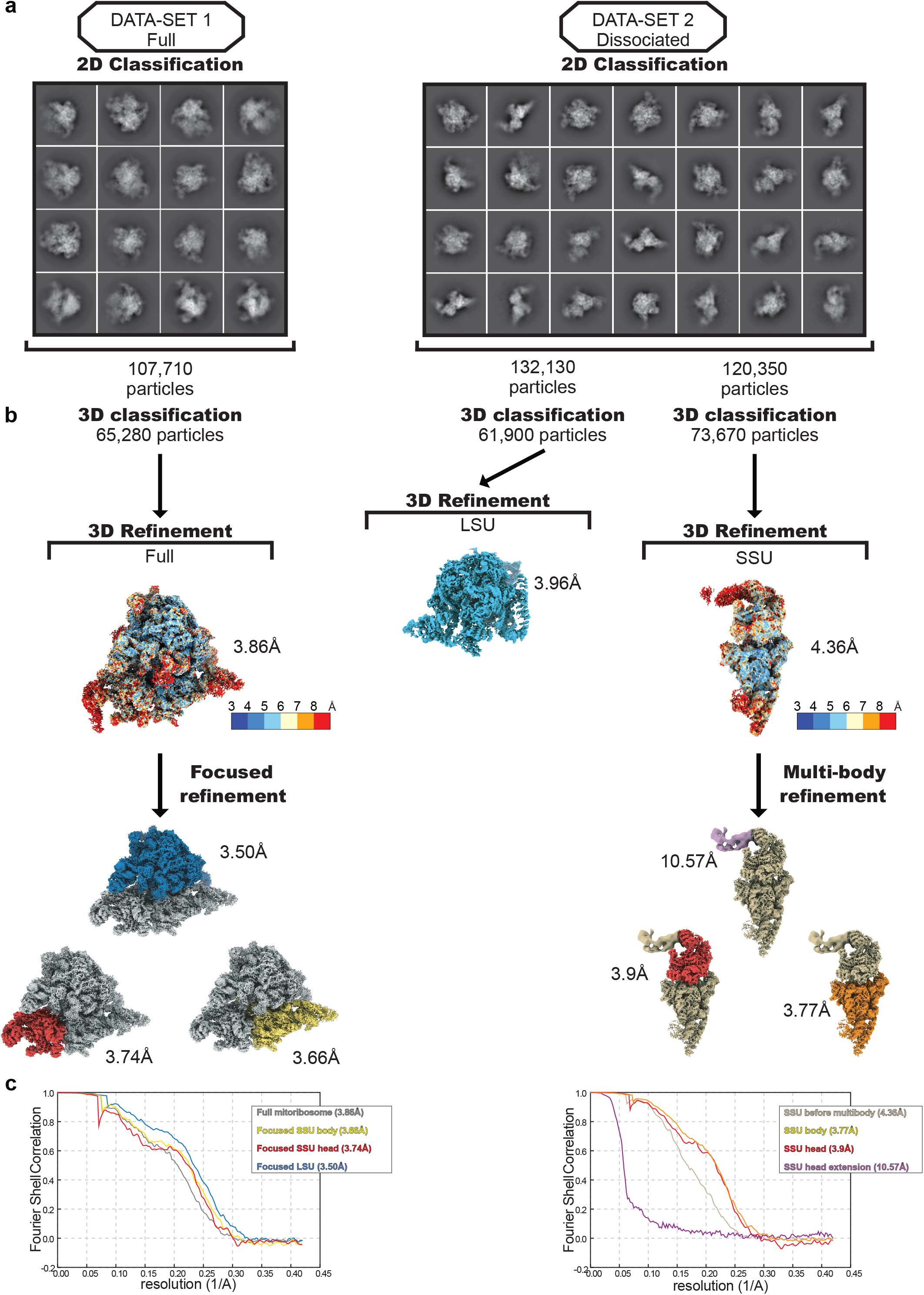
Data processing workflow. Graphical summary of the processing workflow described in Methods, with 2D classes presented in **a** for both datasets and 3D processing, presented in **b**, with ResMap of the full mitoribosome and SSU only before further processing. **c** FSC curves of the full mitoribosome and SSU before and after focused classification and multibody refinement.

**Extended Data Fig. 2.**
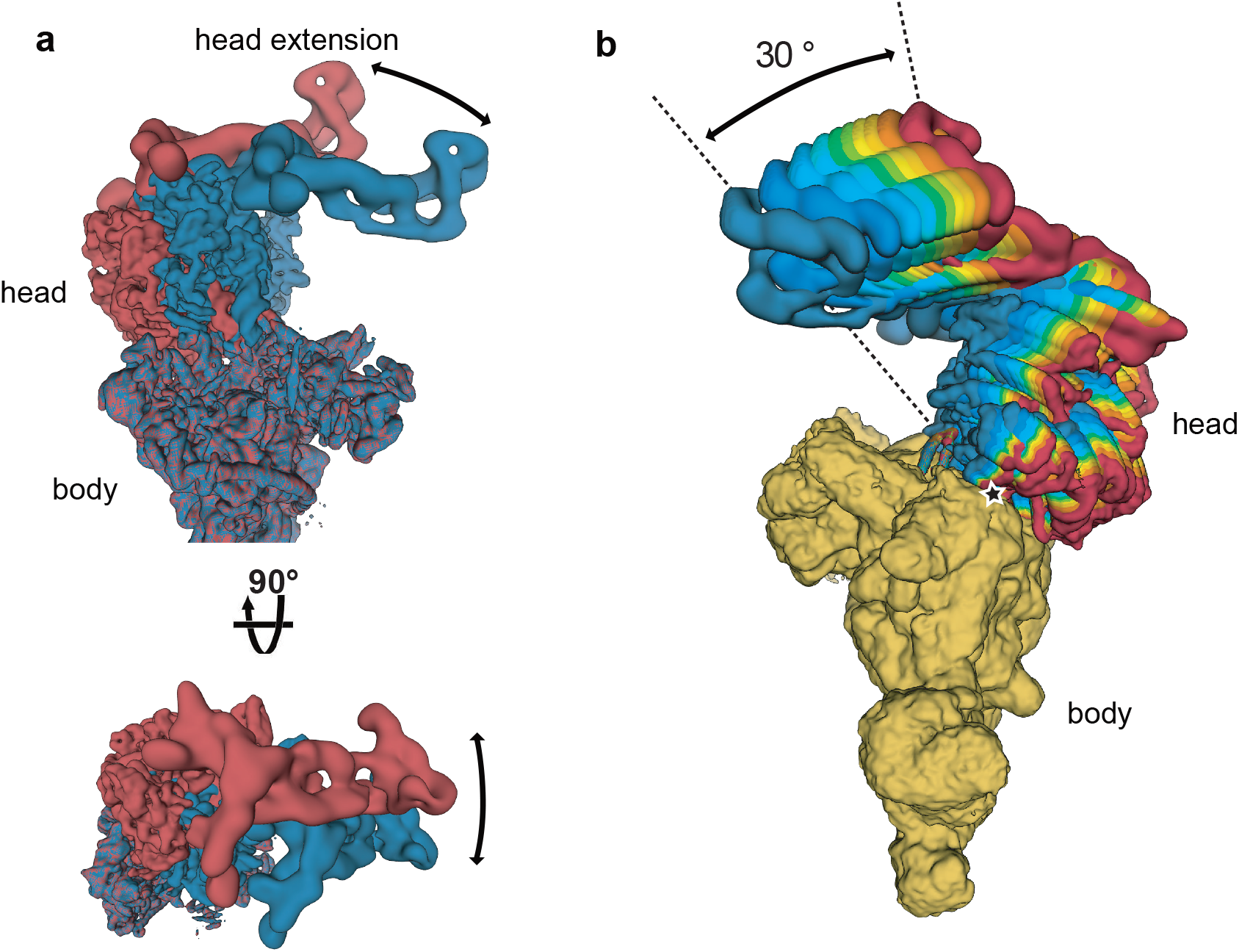
Multibody refinement, additional SSU head domain movement amplitude. **a** Views of the two extreme states of the head and head extension, relative to the body of the SSU, calculated using the multibody refinement implemented in RELION3^12^, showing the movement in two different planes. **b** All ten states reveals a movement amplitude of 30°.

**Extended Data Fig. 3.**
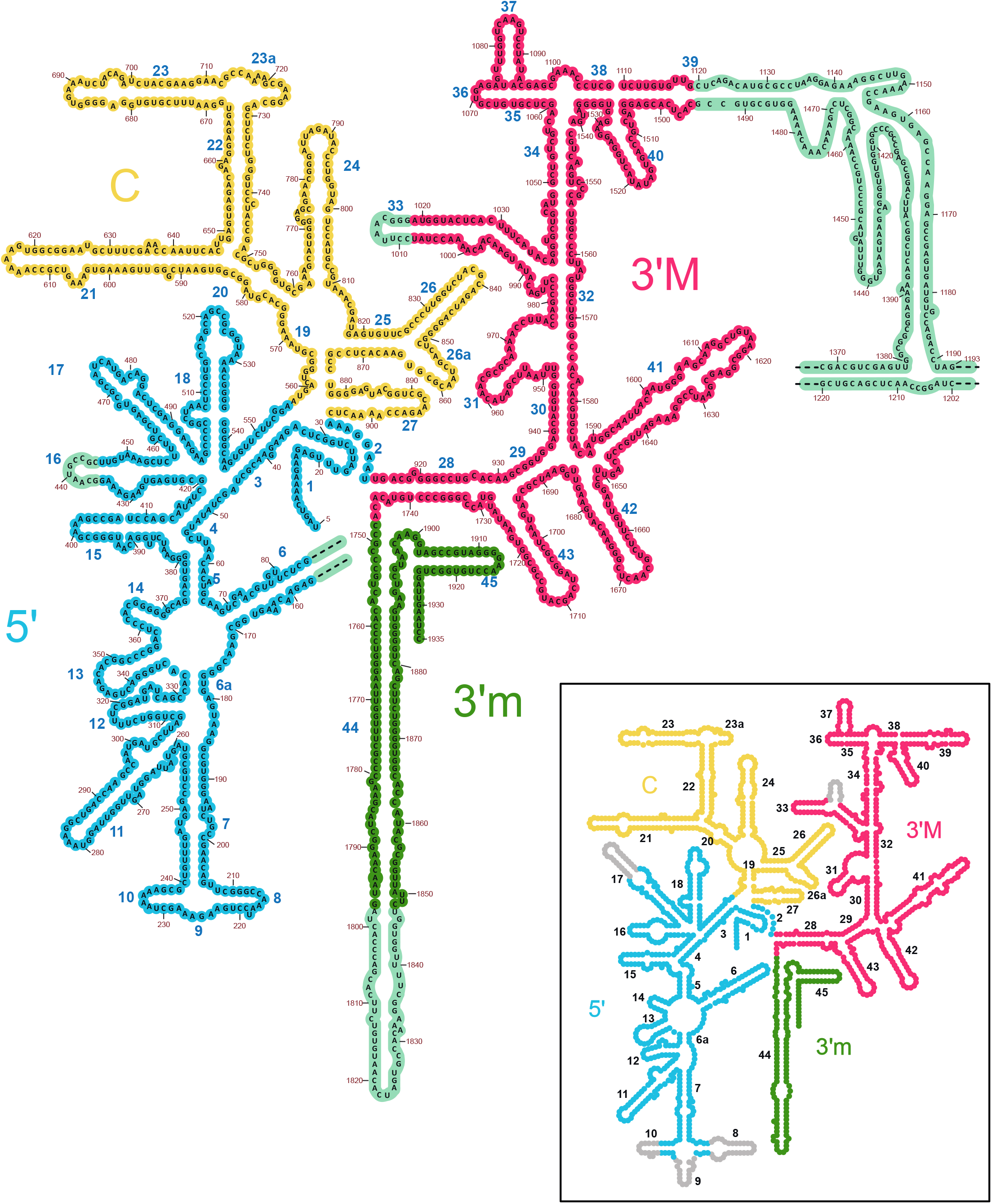
Secondary structure diagram of the plant 18S rRNA. 2D representation of the 18S rRNA colored by domain. The rRNA expansions specific to the plant mitoribosome are highlighted in cyan. Extensions that could not be modelled are indicated by dashed lines. A simplified secondary structure diagram of the E. coli 16S rRNA is also shown in the black frame, helices not present in the plant mitoribosome are shown in gray. Secondary structure templates were obtained from the RiboVision suite (http://apollo.chemistry.gatech.edu/RiboVision)

**Extended Data Fig. 4.**
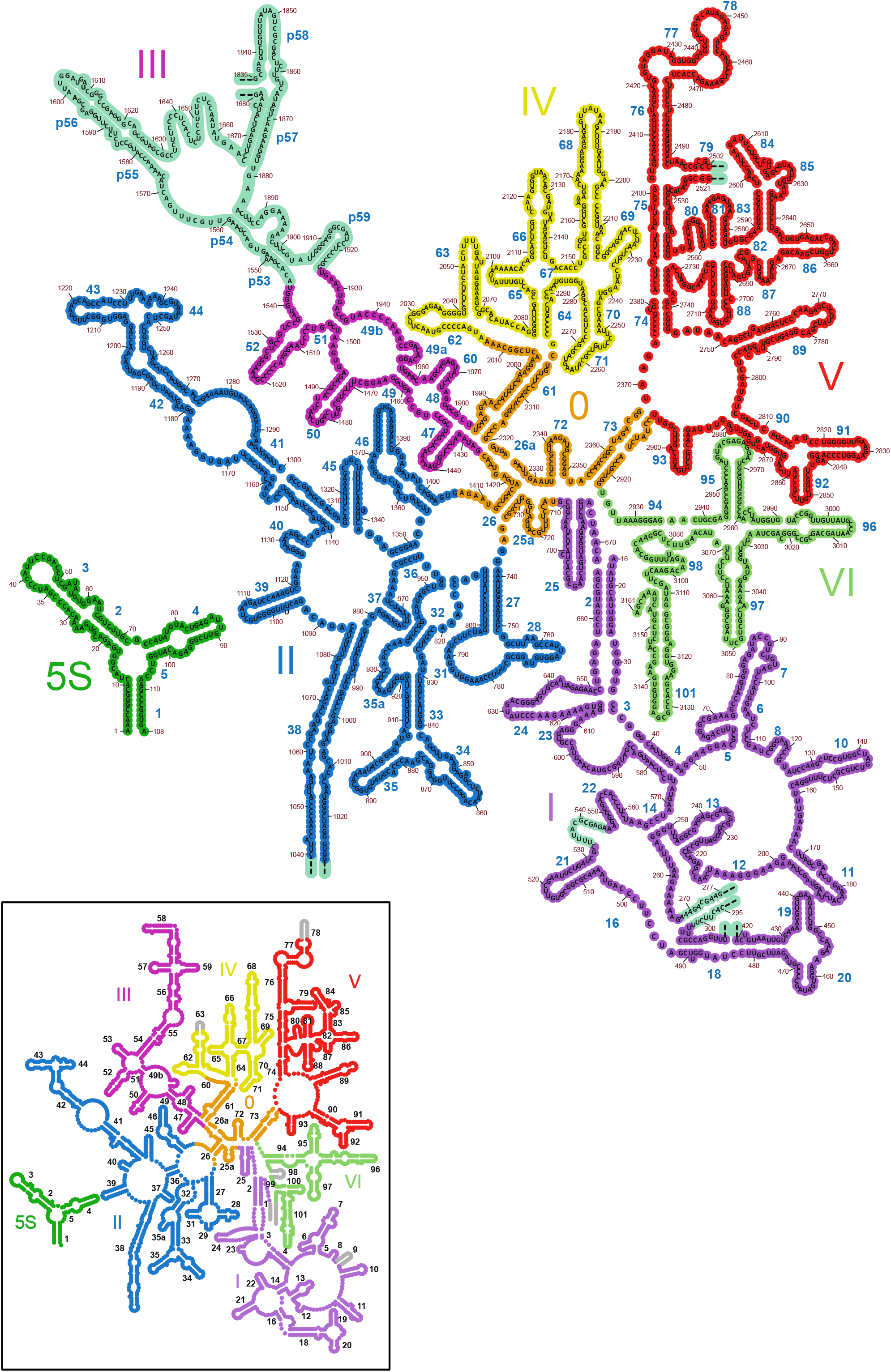
Secondary structure diagram of the plant 26S and 5S rRNA. 2D representation of the 26S rRNA colored by domain. 5S rRNA is shown in dark green. The rRNA expansions specific to the plant mitoribosome are highlighted in cyan. Extensions that could not be modelled are indicated by dashed lines. Simplified secondary structure diagram of the E. coli 23S rRNA is also shown in the black frame, helices not present in the plant mitoribosome are shown in gray. Secondary structure templates were obtained from the RiboVision suite (http://apollo.chemistry.gatech.edu/RiboVision)

**Extended Data Fig. 5.**
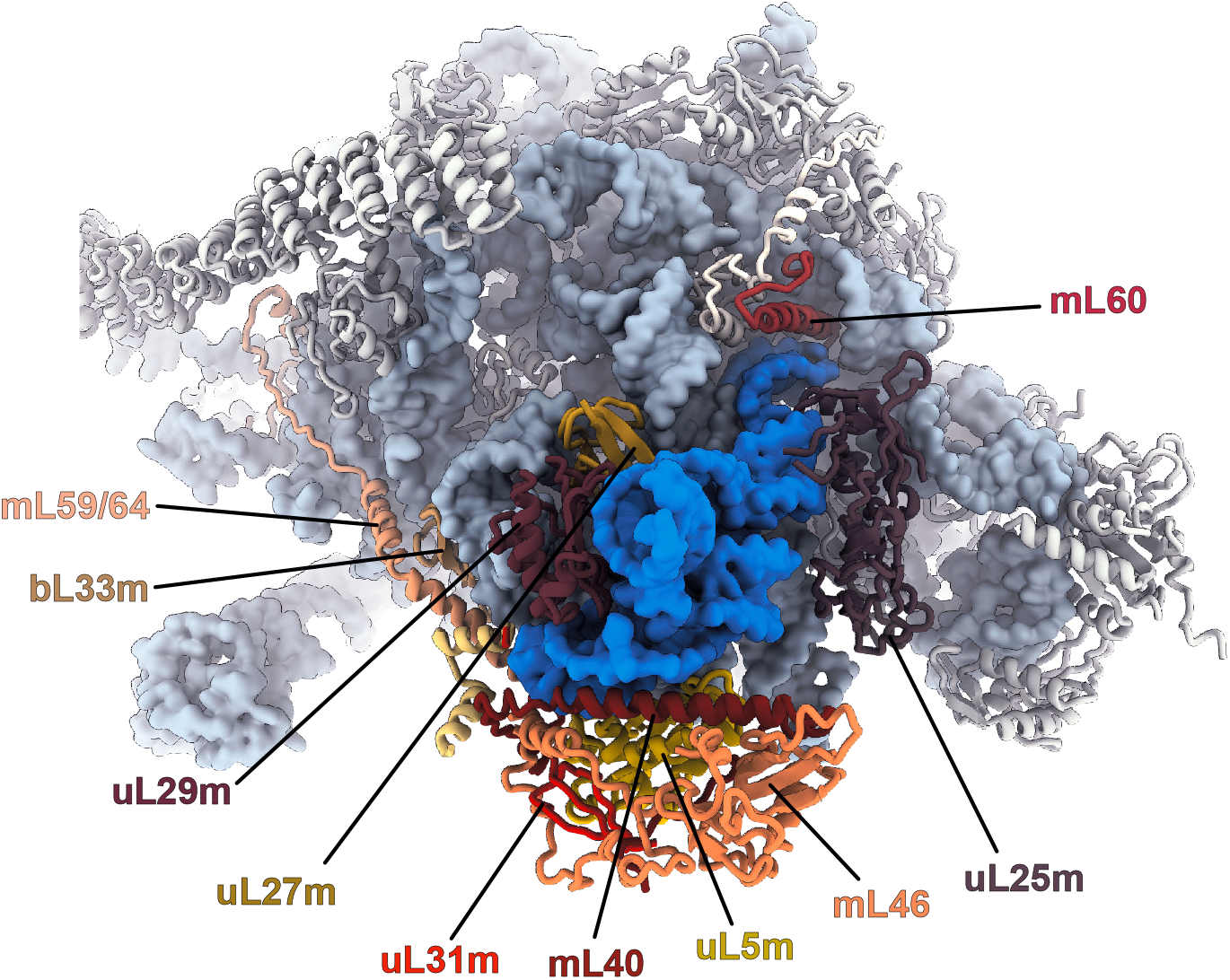
The central protuberance of the plant mitoribosome contains a 5S rRNA. Atomic model of the central protuberance of the plant mitoribosome. Protein components are each indicated in different colors and the 5S rRNA is shown in blue. The plant mitoribosome, contrary to yeast, mammalian and trypanosoma mitoribosome has a 5S rRNA in its CP, like in bacteria. However, the mitochondria specific proteins mL40, mL46, mL59/64 and mL60 are present, augmenting the overall volume of the CP. Moreover, this indicates that the loss of the 5S in yeast, mammals and trypanosoma occurred after the acquisition of these proteins, which seem to constitute core components of the ancestral mitoribosome.

**Extended Data Fig. 6.**
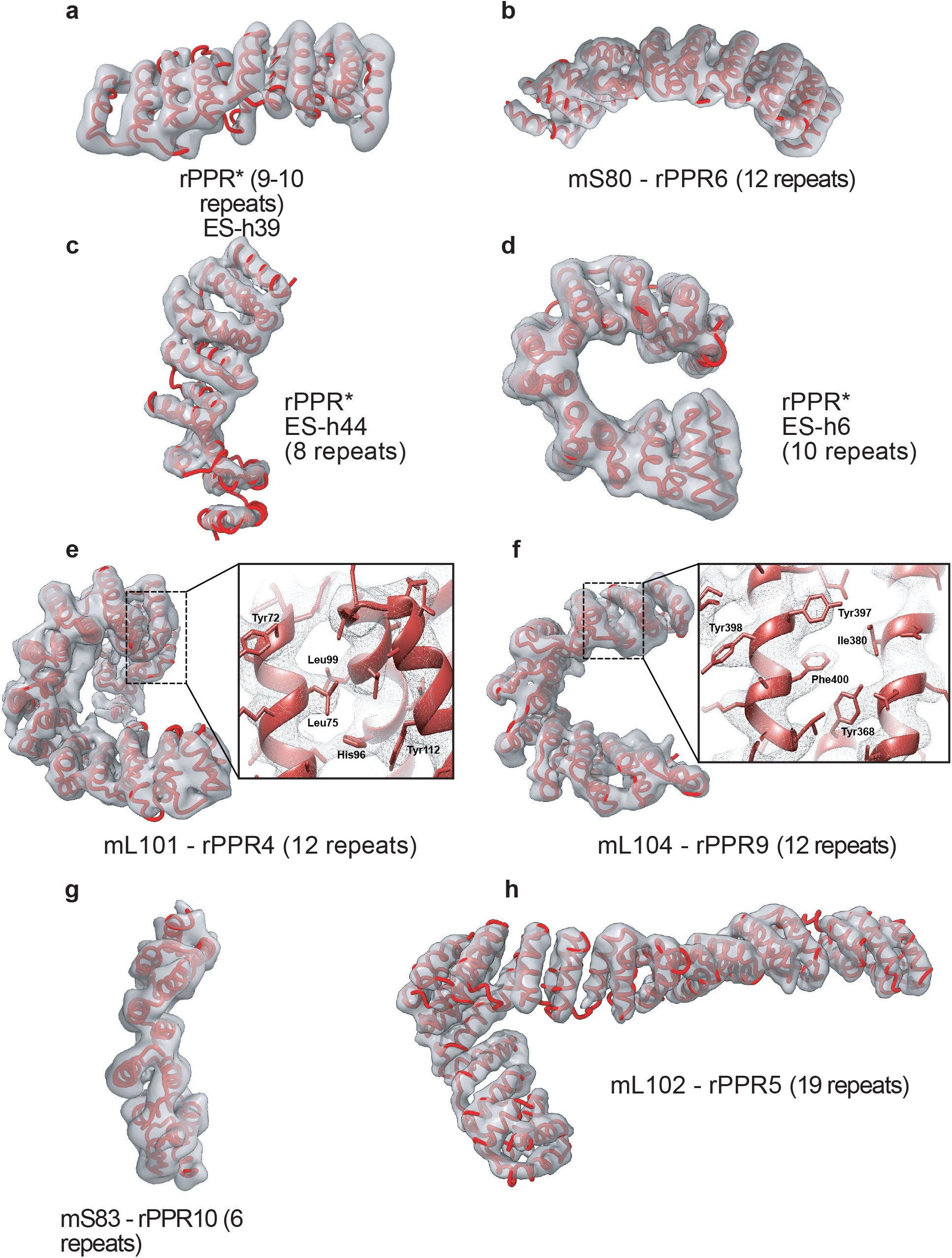
rPPR proteins in their respective densities. **a-h** All rPPR present in the model in their respective filtered densities. Assignment of rPPRs was made mainly thanks to their numbers of repeats that constitute reliable “finger prints”. rPPR 4, 6 and 9 share the same number of repeats, in which case their assignment was possible thanks to the analysis of their bulky side-chains. **a**, **c** and **d** are designated rPPR* as the local resolution could not allow to distinguished them, even based on the number of repeats as rPPR1, 3a and 3b all are predicted to have 10 repeats. **e-f** Detailed with of selected side-chain in their densities that allowed unambiguous assignment. **g** mS83, even though resolved at low resolution, it was identified based on its unique number of repeats (6), which correlates with the predicted number of repeats from the TPRpred software^23^.

**Extended Data Fig. 7.**
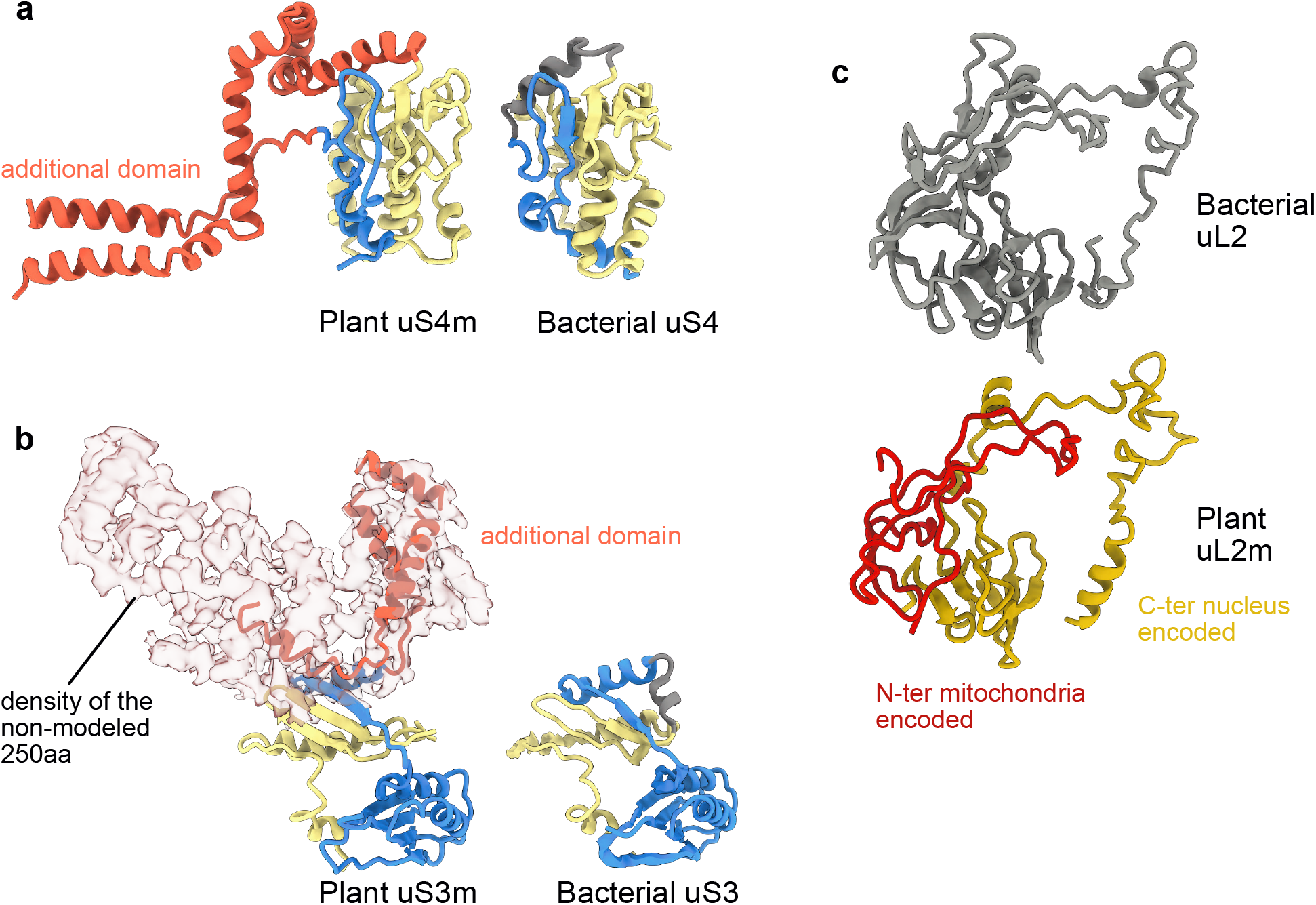
Specificities of plant mitoribosome proteins. **a** Compared view of uS4m and bacterial uS4. The N-terminal part is shown in yellow and the C-terminal part in blue. The additional domain of uS4m is shown in red. This additional domain makes a part of the SSU body protuberance. **b** Similar to uS4m, uS3m also present a large additional domain. Due to low resolution in this area only part of the insertion was modelized, however there is no doubt that the 250 amino-acids missing would constitute the large density observed on the head of the SSU, shown here in brown. N-terminal parts of the proteins are shown in yellow and C-terminal parts in blue. The additional domain of uS3m is shown in red. **c** In plant mitochondria, the uL2m protein was already speculated to be composed of two parts^24^. The structure confirmed this hypothesis. The N-terminal part of the protein (red) is encoded by a mitochondrial gene and the C-terminal part of the protein (yellow) is encoded by a nuclear gene.

**Extended Data Fig. 8.**
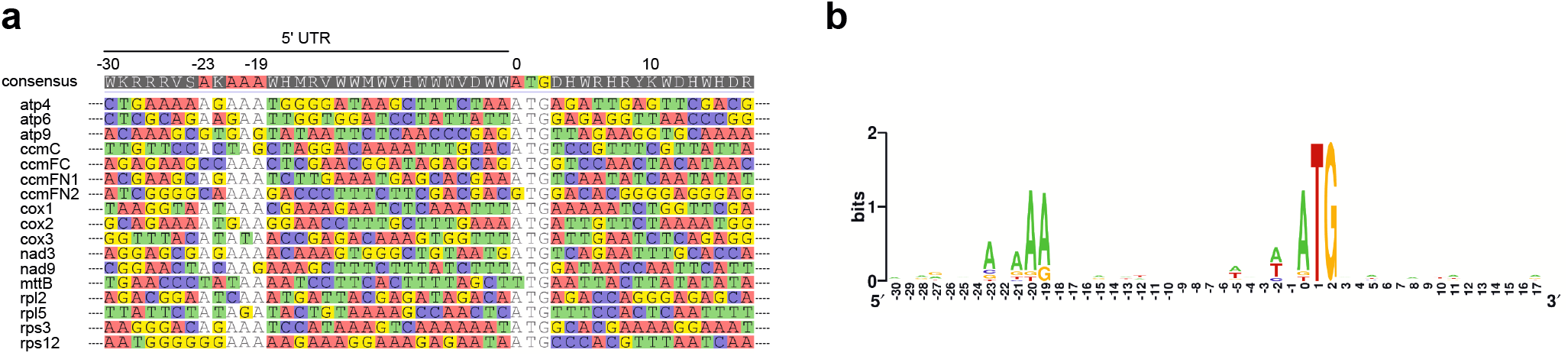
5’UTR of the mitochondrial mRNAs. **a** Alignment of sequences surrounding the initiation codons of the 17, out the 33, protein coding genes encoded in the Arabidopsis mitochondrial genome. A characteristic AxAAA consensus is observed and illustrated by the WebLogo (https://weblogo.berkeley.edu/logo.cgi) representation in **b.**

